# Chronic intermittent ethanol exposure selectively increases synaptic excitability in the ventral domain of the rat hippocampus

**DOI:** 10.1101/337097

**Authors:** Sarah E. Ewin, James W. Morgan, Farr Niere, Nate P. McMullen, Samuel H. Barth, Antoine G. Almonte, Kimberly F. Raab-Graham, Jeffrey L. Weiner

**Affiliations:** Department of Physiology and Pharmacology Wake Forest School of Medicine Winston-Salem, NC 27157

**Author notes:** Corresponding author: Sarah E. Ewin Department of Physiology & Pharmacology Wake Forest School of Medicine, Winston-Salem, NC 27157 USA.

**Keywords:** Chronic intermittent ethanol, ventral hippocampus, anxiety

## Abstract

Many studies have implicated hippocampal dysregulation in the pathophysiology of alcohol use disorder (AUD). However, over the past twenty years, a growing body of evidence has revealed distinct functional roles of the dorsal (dHC) and ventral (vHC) hippocampal subregions, with the dHC being primarily involved in spatial learning and memory and the vHC regulating anxiety-and depressive-like behaviors. Notably, to our knowledge, no rodent studies have examined the effects of chronic ethanol exposure on synaptic transmission along the dorsal/ventral axis. To that end, we examined the effects of the chronic intermittent ethanol vapor exposure (CIE) model of AUD on dHC and vHC synaptic excitability. Adult male Long-Evans rats were exposed to CIE or air for 10 days (12 hrs/day; targeting blood ethanol levels of 175-225 mg%) and recordings were made 24 hours into withdrawal. As expected, this protocol increased anxiety-like behaviors on the elevated plus-maze. Extracellular recordings revealed marked CIE-associated increases in synaptic excitation in the CA1 region that were exclusively restricted to the ventral domain of the hippocampus. Western blot analysis of synaptoneurosomal fractions revealed that the expression of two proteins that regulate synaptic strength, GluA2 and SK2, was dysregulated in the vHC, but not the dHC, following CIE. Together, these findings suggest that the ventral CA1 region may be particularly sensitive to the maladaptive effects of chronic ethanol exposure and provide new insight into some of the neural substrates that may contribute to the negative affective state that develops during withdrawal.

**Highlights:**

Chronic intermittent ethanol exposure produces robust increases in anxiety-like behavior in male Long Evans rats.
Chronic intermittent ethanol exposure increases synaptic excitability in the ventral, but not the dorsal, domain of the hippocampus.
These changes in excitability are associated with alterations in synaptoneurosomal expression of small conductance calcium-activated potassium channels and the GluA2 AMPA receptor subunit that are also restricted to the ventral hippocampus.

## 1. Introduction

Alcohol use disorder (AUD) affects over 16.3 million Americans and is a worldwide socioeconomic and public health problem, accounting for more than 6% of the global burden of disease (Center for Behavioral Health Statistics, 2015; World Health Organization, 2003). AUD is a chronic relapsing disease that is characterized by a transition from recreational drinking to excessive alcohol use, involving a shift from positive reinforcement to negative reinforcement (Koob, 2015; Koob and Volkow, 2016). In other words, individuals are initially motivated to drink primarily because of the pleasurable effects of alcohol consumption but eventually become drawn to alcohol to avoid the negative affective feelings that emerge during periods of abstinence.

Many studies have shown that anxiety represents an important element of the negative affective state that develops during withdrawal from chronic alcohol exposure (Becker, 2008; Breese et al., 2011). In fact, individuals with anxiety and stressor-related disorders are 2-4 times more likely to develop AUD, are diagnosed with AUD significantly earlier than individuals without comorbid anxiety disorders, and this dual diagnosis is associated with much poorer treatment outcomes (Kushner et al., 2011, 2005; Smith and Randall, 2012). Withdrawal-associated anxiety is also a major contributing factor to relapse in treatment–seeking individuals (Sinha et al.2011). Additionally, stress and alcohol-associated cues trigger robust increases in craving in abstinent alcoholics that are associated with increased anxiety and negative affect and the magnitude or intensity of this craving is a strong predictor of relapse (Sinha et al., 2009).

Despite the clinical importance of understanding the neural substrates responsible for the negative affective state that develops following chronic alcohol exposure and withdrawal, much remains unknown regarding the specific brain regions and circuits that mediate withdrawal-associated increases in anxiety behaviors and alcohol drinking. One preclinical model of AUD that has been extensively validated to address these questions is the chronic intermittent ethanol (CIE) vapor exposure regimen (Becker, 2017; Gilpin et al., 2008; Reynolds and Berridge, n.d.; Vendruscolo and Roberts, 2014). Using this model, which involves daily cycles of ethanol vapor exposure followed by a withdrawal period, rats and mice develop physiological and behavioral signs of ethanol dependence, including marked escalations in ethanol self-administration (Criado and Ehlers, 2013; Finn et al., 2007; O’Dell et al., 2004) and increases in anxiety-like behaviors on assays like the elevated plus-maze (Cagetti et al., 2004).

Elegant neurobiological studies used the CIE procedure to identify key elements of the neural circuitry that drives the maladaptive withdrawal-associated behaviors promoted by this model. Most recent studies have focused on the prefrontal cortex, amygdala nuclei, and the bed nucleus of the stria terminalis, regions that comprise interconnected circuits known to play an integral role in negative affective states. These studies have shown that withdrawal from CIE increases synaptic excitability within these circuits and that these maladaptive changes drive negative affective behaviors (Christian et al., 2012b; Conrad and Winder, 2011; de Guglielmo et al., 2016; Den Hartog et al., 2016; Diaz et al., 2011; Holmes et al., 2012; Läck et al., 2007; Marcinkiewcz et al., 2016; Pleil et al., 2015).

The hippocampus is another brain region that is intimately connected to the circuitry that governs negative emotional states. This structure runs along a ventral-dorsal axis in rodents, which corresponds to an anterior-posterior axis in humans (Strange et al., 2014). While the intrinsic circuitry of the dorsal and ventral domains is similar, these subregions are comprised of distinct afferent and efferent projections (Strange et al., 2014). Notably, the ventral domain of the hippocampus makes strong monosynaptic, reciprocal connections to several nodes of the emotional network and has long been known to play an integral role in anxiety-like behaviors (Bannerman et al., 2003; Fanselow and Dong, 2010; Felix-Ortiz et al., 2013; Huff et al., 2015; Kjelstrup et al., 2002; Maggio and Segal, 2009, 2007; Strange et al., 2014). In support of this, recent optogenetic studies have demonstrated that the excitatory projection from the basolateral amygdala (BLA) to the ventral hippocampus (vHC) can bidirectionally modulate anxiety-like behaviors in a manner similar to that observed via manipulations of the canonical basolateral-central amygdala “anxiety” circuit (Felix-Ortiz et al., 2013; Felix-Ortiz and Tye, 2014). Moreover, optical inhibition of the BLA-vHC circuit also disrupts the consolidation of footshock-associated fear but not contextual fear learning (Huff et al., 2015). In contrast, the dHC contains the greatest density of place cells that encode spatial location and these cells send strong excitatory projections to areas like the dorsal subiculum, retrosplenial cortex and anterior cingulate cortex, regions known to play an integral role in cognitive processing of visual information (Fanselow and Dong, 2010; Jung et al., 1994; Kjelstrup et al., 2002; Potvin et al., 2007; Tannenholz et al., 2014).

Although several studies have reported that CIE promotes increases in hippocampal excitability (Läck et al., 2007; Nelson et al., 2005; Roberto et al., 2001), to our knowledge none have directly compared the effects of CIE on synaptic transmission in the dorsal and ventral domains of the hippocampus. To that end, we employed electrophysiological and biochemical approaches to assess the effects of CIE on synaptic excitability in the rat dHC and vHC. We hypothesized that withdrawal following CIE would promote increases in synaptic excitability in both subregions of the hippocampus. However, given that negative affective behaviors are some of the most sensitive alterations observed following ethanol withdrawal (Morales et al., 2018; Rose et al., 2016; Sidhu et al., 2018), we also predicted that CIE-associated synaptic alterations would be most robust in the vHC. Here we report that withdrawal following CIE markedly increases measures of synaptic excitability in the ventral domain of the hippocampus while modestly decreasing synaptic excitation in the dorsal subregion. We further identify changes in two synaptic proteins, the AMPA receptor (AMPAR) subunit GluA2 and the small conductance Ca^2+^-activated K^+^-channel subunit SK2, that may contribute to these divergent adaptations.

## 2. Methods

### 2.1 Subjects

Male Long Evans rats were purchased from Envigo, IN and arrived at 175-200g. Upon arrival rats were singly housed in clear cages (25.4 cm × 45.7cm) and maintained on a reverse 12 hour: 12 hour light dark cycle with lights on at 9pm. Rats had *ad libitum* access to food (Prolab RMH 3000, LabDiet: PMI Nutrition International, St. Louis, MO) and water throughout the study. Animal care procedures were carried out in accordance with the NIH Guide for the Care and Use of Laboratory Animals and were approved by the Wake Forest University Animal Care and Use Committee.

### 2.2 Chronic intermittent ethanol vapor inhalation exposure

Animals in the CIE condition were housed in their standard home cages which were placed in custom-built Plexiglas chambers (Triad Plastics, Winston-Salem, NC). Ethanol vapor was pumped into the chamber for 12 hours a day for 10 consecutive days during the light cycle (9pm to 9am). Control animals (AIR) were housed in the same manner with their home cages placed inside the custom-built Plexiglas chambers on the same light cycle, but were exposed only to room air. Animals were weighed daily and tail blood samples were taken every other day during the 10 day CIE procedure at 9am to monitor blood ethanol concentrations (BECs). Following the 10 days of CIE, animals underwent 24 hours of withdrawal (no ethanol vapor) and were then run on behavioral assay or sacrificed for electrophysiological or biochemical studies.

### 2.3 Blood Ethanol Determination

Blood ethanol concentrations (BECs) were measured every other day at 9am. 10µL of blood was collected from a tail snip of each rat. BECs were determined using a commercially available alcohol dehydrogenase enzymatic assay kit (Carolina Liquid Chemistries Corporation, Brea, CA). Ethanol concentrations were then determined using a spectrophotometer (Molecular Devices Spectra Max). The target range for BECs was 175 – 225 mg/dL.

### 2.4 Elevated Plus-Maze

Anxiety-like behavior was assessed in a subset of the CIE and AIR animals following 24 hours of withdrawal using standard elevated plus-mazes (Med Associates, St. Albans, VT) raised 72.4 cm from floor level, with runways measuring 10.2 cm wide by 50.8 cm long. Open runways had 1.3 cm high lips and closed runways were enclosed in 40.6 cm high black polypropylene walls. Exits and entries from runways were detected via infrared sensors attached to the opening of each arm of the maze. Data were obtained and recorded via personal computer interfaced with control units and MED-PC programming (Med Associates). Animals were placed at the junction of the four arms at the beginning of the session, and activity was measured for five minutes. Anxiety-like behavior was assessed by measuring the total time spent on the open arms of the maze as well as the number of entries into the open arms. General locomotor activity was assessed by measuring the number of closed arm entries.

### 2.5 Open Field Test

Immediately following the elevated plus maze, general locomotion in a novel environment was measured in all animals using an open field test conducted in Plexiglass chambers (41.5 cm ×41.5cm ×30 cm). At the start of the test, animals were placed in the center of the chambers equipped with Omnitech Superflex Sensors (Omnitech Electronics, Inc.), which utilize arrays of infrared photodetectors located at regular intervals along each way of the chambers. The chamber walls were solid and contained within sound attenuating boxes with 15 watt light bulbs to illuminate the arena. Exploratory activity in this environment was measured for 30 minutes, and data were analyzed in five minute time bins.

### 2.6 Electrophysiology

After the induction of a deep anesthetic plane with isoflurane, rats were decapitated, and their brains removed and placed in ice-cold cutting artificial cerebral spinal fluid (aCSF) consisting of 85mM NaCl, 1.25 mM NaH_2_PO_4_, 25 mM NaHCO_3_, 10 mM D-Glucose, 75 mM sucrose, 3 mM KCl, 7mM MgCl_2_, 0.5 mM CaCl_2_, and 0.6 mM ascorbate bubbled with 95% O_2_ and 5% CO_2_. Transverse slices containing the dHC and vHC were cut at a thickness of 375μm using a VT1000S Vibratome (Leica Microsystems)., Differentiation of ventral and dorsal slices was noted prior to recording using definitions from Maruki et al (2001) and Fanselow and Dong (2010). Incubation of slices occurred for at least one hour at room temperature (21-23 °C) in aCSF consisting of 125 mM NaCl, 1.25 mM NaH_2_PO_4_, 25 mM NaHCO_3_, 10 mM D-Glucose, 2.5 mM KCl, 1 mM MgCl_2_, and 2 mM CaCl_2_ bubbled with 95% O_2_ and 5% CO_2_ before experiments commence. Slices were transferred to a recording chamber and perfused with oxygenated, heated (to 32 °C) aCSF at 2mL/min. Filamented borosilicate glass capillary tubes (inner diameter, 0.86 μm) were pulled using a horizontal pipette puller (P-97; Sutter Instrument) to prepare recording electrodes (1-3 MΩ resistance). To acquire extracellular field potentials the glass capillary tubes were filled with 0.9% saline. Extracellular field recordings were obtained from Schaffer collateral → CA1 synapses of the dHC or vHC using nickel dichromate bipolar stimulating electrodes. For input-output curves, field excitatory post synaptic potentials (fEPSP) were evoked every 10 seconds at 10, 20, 50, 100, 150, 200, 300, 500, and 700 µA five times per stimulus intensity. All recordings were acquired using an Axoclamp 2B amplifier, digitized (Digidata 1321A; Molecular Devices) and analyzed with pClamp 10.4 software (Molecular Devices).

### 2.7 Western blotting

Western blot analyses were performed on synaptoneurosomes (SNs) that were obtained from hippocampal slices that were prepared similarly for electrophysiology experiments. (See Electrophysiology methods above). Briefly, slices were homogenized in buffer (50 mM Tris, pH 7.35; protease and phosphatase inhbitors (Halt, ThermoFisher)). Homogenates were sequentially filtered through 100 μm and 5 μm filters to produce SNs(Niere et al., 2016; Quinlan et al., 1999; Workman et al., 2013). SNs were centrifuged (14,000g, 20 min, 4°C) to obtain a pellet that was solubilized in RIPA buffer (150 mM NaCl; 10 mM Tris, pH 7.4; 0.1% SDS; 1% Triton X-100; 1% deoxychoate 5 mM EDTA; Halt). The insoluble fraction of SNs was removed by centrifugation at 14,000g, 20 min, 4°C. The soluble fraction was used for immunoblot analysis. 50 μg of protein were run for each sample and separated by SDS-PAGE. The following antibodies were used to visualize the proteins of interest: mouse anti-GluA2 (1:2000; Neuromab; Davis, CA); rabbit anti-SK2 (1:1000; Alomone Lab, Jerusalem, Israel); mouse anti-actin (1:10,000; Sigma; St. Louis, MO). To visualize the proteins, membranes were incubated in fluorescence-conjugated secondary antibodies (AF680; AF800; 1:4000; LiCor, Lincoln, NE) and imaged using the Odyssey CLx infrared imaging system. For densitometry analysis of proteins, ImageJ software (National Institutes of Health) was used.

### 2.8 Data Analysis and Statistics

Elevated plus-maze and Western blot data were analyzed using unpaired t-tests, or Mann-Whitney Rank Sum Tests in the event of non-normally distributed data. Open field test data and electrophysiology data were analyzed using two way repeated measures ANOVAs. Where noted, *post hoc* analysis was conducted using Bonferroni’s multiple comparison test. A generalized linear model was run in SAS to examine the relationship between CIE, fiber volley amplitude and fEPSP slope using ANCOVAs. The minimal level of significance was set as *p*< 0.05 for all analyses.

## 3. Results

### 3.1 Chronic intermittent ethanol exposure increases anxiety-like behavior

Anxiety-like behavior was assessed 24 hr following the CIE exposure paradigm using the elevated plus-maze. CIE animals (n= 12) exhibited an increases in anxiety-like behavior, as evidenced by less time spent on the open arms (Figure 1A; t(22) = 198.5; p = 0.005; two-tailed) and less open arm entries (Figure 1B; t(22) = 198.0; p = 0.005; two-tailed). CIE and AIR animals showed a significant difference in closed arm entries (Figure 1C; t(22)= 4.168; p <0.001), typically used as a measure of non-specific locomotion. However, CIE animals did not differ from AIR animals in locomotor activity in response to a novel environment (Figure 1D), as evidenced by no significant main effect of condition (AIR or CIE) (F_1, 110_ = 0.128; p =0.72). There was a significant main effect of time (F_1, 110_ = 61.155; p<0.001), as rodents acclimated to the novel arena, but no significant interaction between condition and time (F_1, 110_ = 0.445; p =0.82).

**Figure 1.**
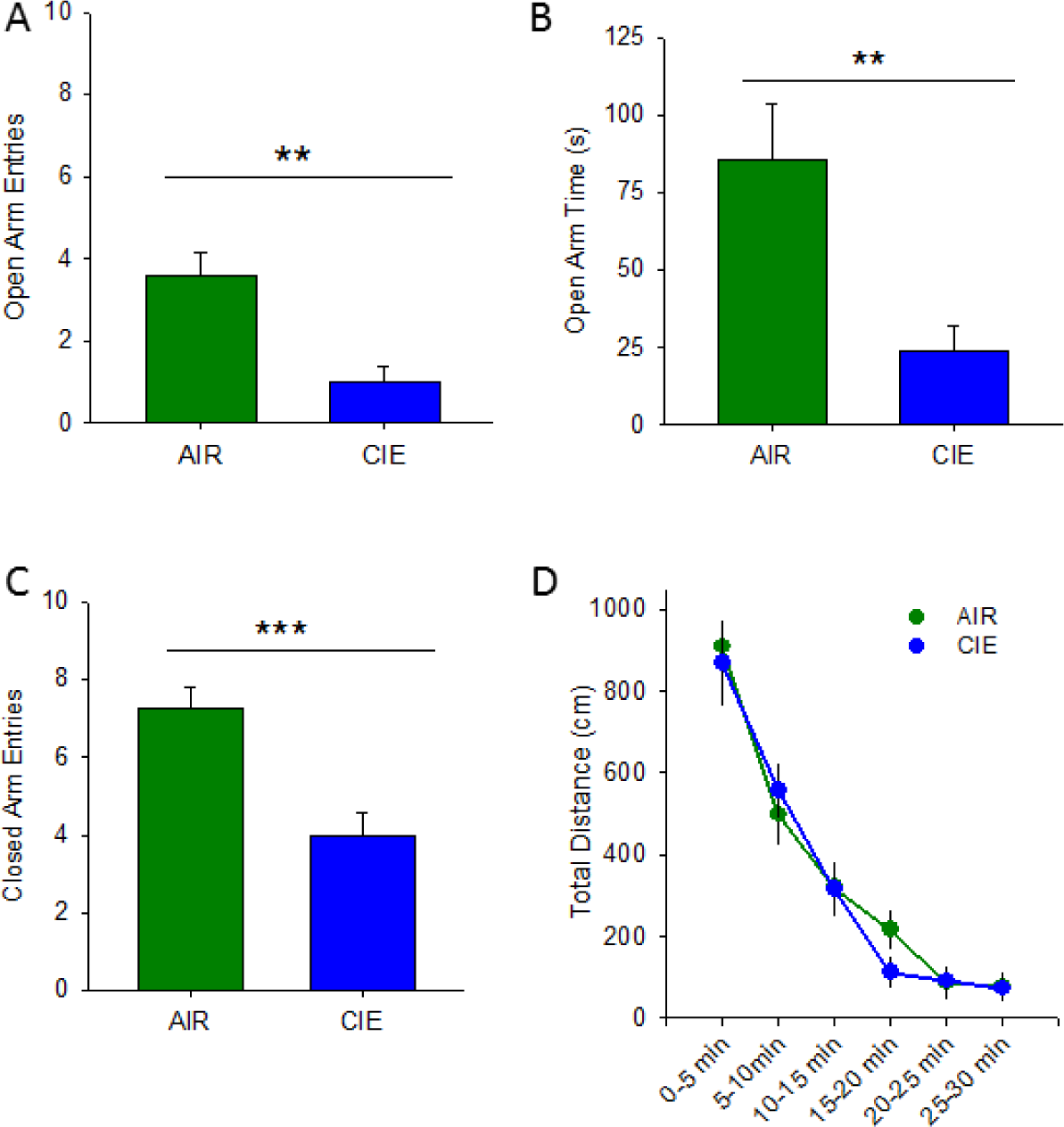
Chronic intermittent ethanol increases anxiety-like behavior in the elevated plus-maze. AIR exposed (N =12) rats exhibited less anxiety-like behavior on the elevated plus-maze than CIE (N =12) counterparts, as evidenced by more time spent in the open arms (A) and more open arm entries (B). AIR rats also exhibited a greater number of closed arm entries (C). Additionally, AIR and CIE rats do not exhibit differences in locomotor activity in the open field test (D; **, p<0.01, ***. p<0.001).

### 3.2 Chronic intermittent ethanol exposure produces divergent effects on hippocampal excitability

To determine whether CIE altered hippocampal excitability, extracellular field excitatory postsynaptic potentials (fEPSPs) were recorded in hippocampal Schaffer-collateral-CA1 region (Almonte et al., 2017). All recordings were conducted at 24-32 hours into withdrawal, a time range similar to that associated with the CIE-associated increase in anxiety-like behavior on the elevated plus-maze. We recorded afferent fiber volley amplitudes and the rising slope of fEPSPs in response to increasing stimulus intensity levels to generate input-output curves. In the vHC (Figure 2A), there was no main effect of treatment condition on fiber volley amplitude (Figure 3A; F_1, 95_ = 0.963; p = 0.35), but there was a main effect of stimulation intensity (F_1, 95_ = 50.796; p <0.001). Although there was no significant interaction between treatment condition and stimulation intensity (F_1, 95_ = 1.472; p = 0.13), given our *a prioiri* hypothesis that CIE would promote increases in synaptic excitability, we conducted additional post-hoc analyses which revealed a significant facilitatory effect of CIE on fiber volley amplitude, albeit only at the highest stimulus intensity tested (Figure 3A; p = 0.01). In regards to the fEPSP slope (Figure 3B), there was a main effect of treatment condition (F_1, 95_ = 7.231; p = 0.02), as well as stimulation intensity (F_1, 95_ = 46.302; p < 0.001). In addition, there was a significant interaction between condition and stimulation intensity on fEPSP slope (F_1, 95_ = 46.302; p <0.001). We then used a model to conduct an analysis of covariance to further understand how withdrawal from CIE altered the relationship between afferent volley amplitude and fEPSP slope. This analysis revealed a significant effect of fiber volley amplitude (Figure 3C; F_1,124_= 108.26; p < 0.0001), but not treatment condition (F_1,124_= 0.01; p = 0.94), on fEPSP slope. There was also a significant interaction between fiber volley amplitude and CIE on fEPSP slope (F_1,124_= 47.01; p <0.0001). This analysis yielded a parameter estimate of 1.039, which indicates that, in the vHC of CIE animals, the fiber volley’s positive effect on fEPSP slope is greatly increased, resulting in an overall enhancement of excitability.

**Figure 2.**
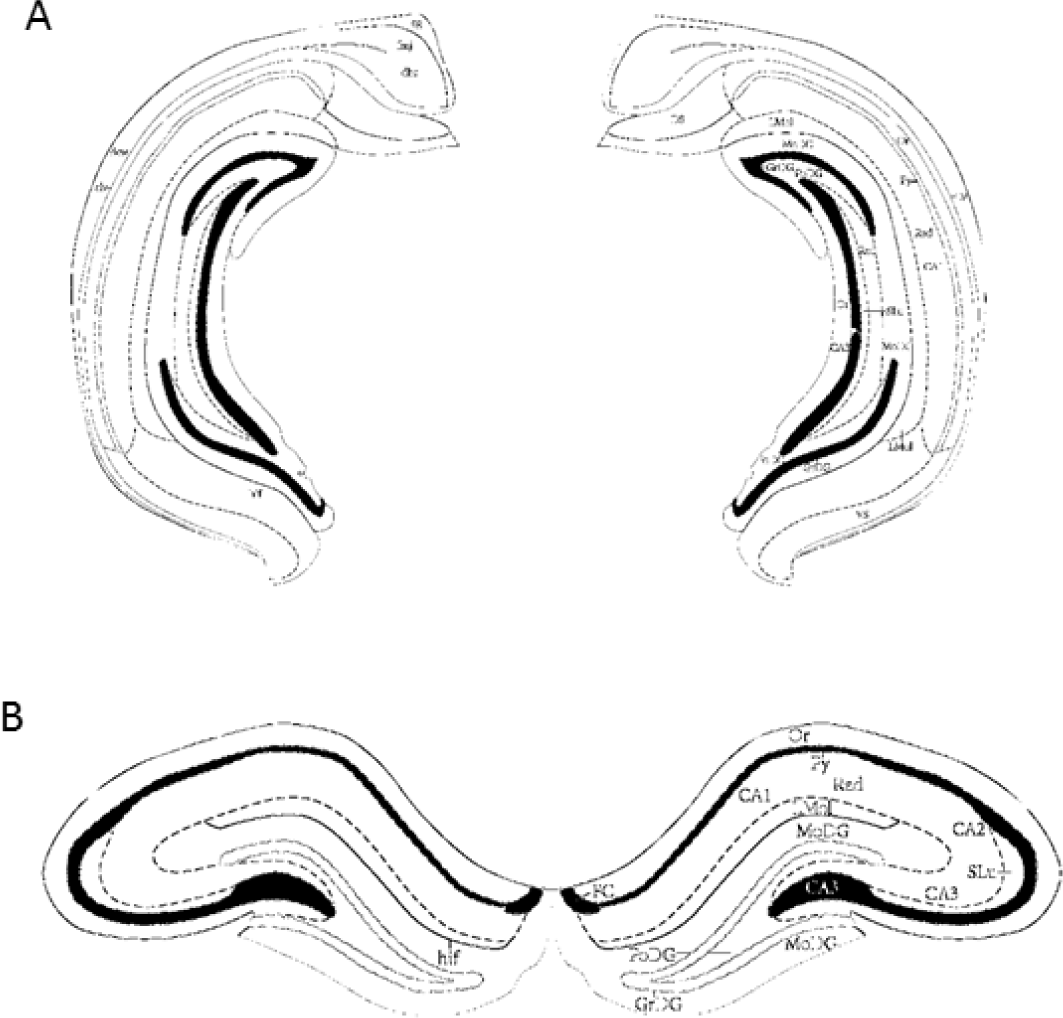
Examples of slices taken from ventral and dorsal hippocampus. Representative images adapted from Paxinos and Watson (2005) depicting the ventral (A; Bregma -5.52mm) and dorsal hippocampus (B; Bregma -3.12mm) for extracellular field recordings.

**Figure 3.**
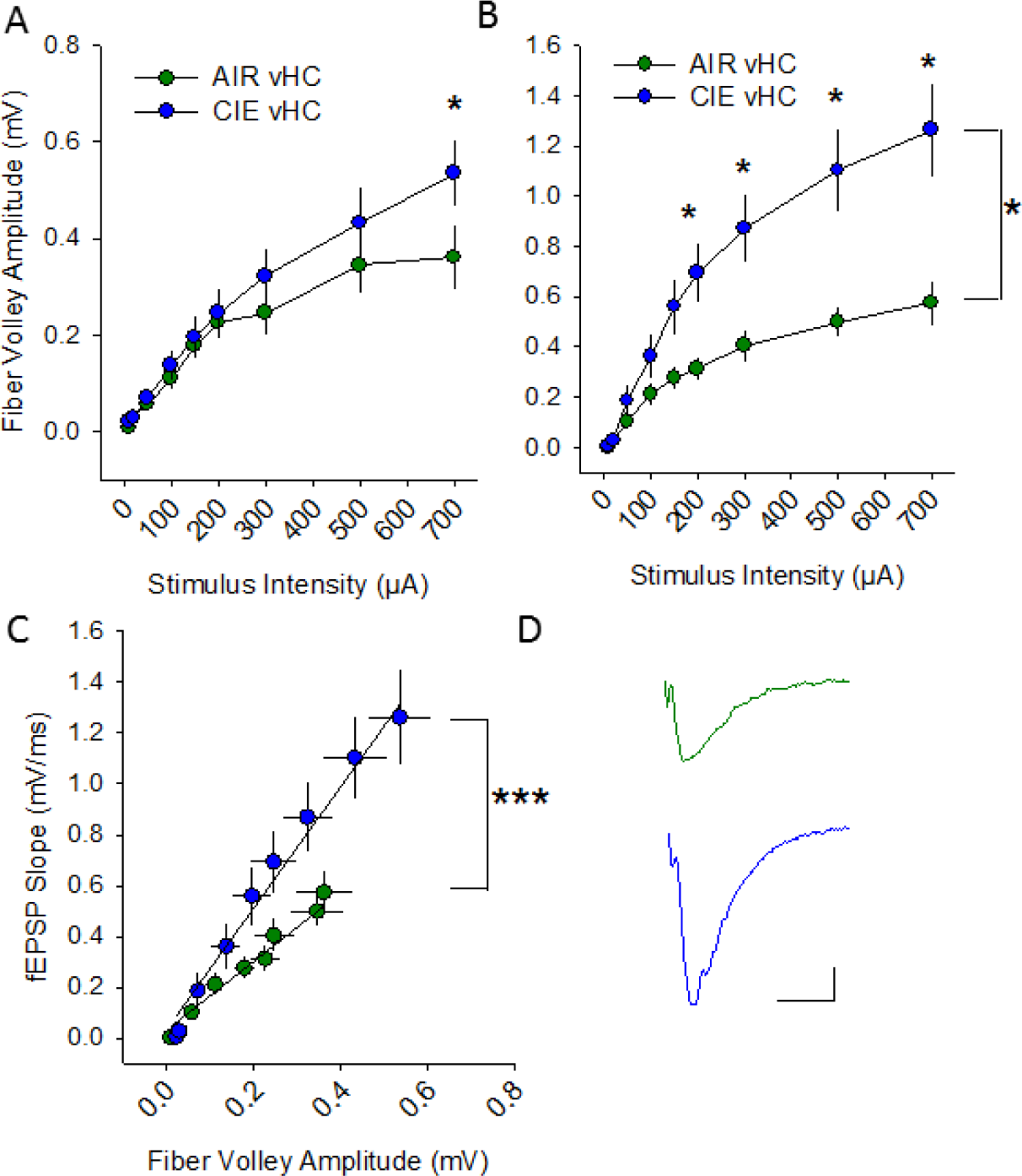
Withdrawal from chronic intermittent ethanol increases synaptic excitability at vHC Schaffer collateral-CA1 synapses. Plots of stimulation intensity vs fiber volley amplitude (A) and fEPSP slope (B) in vHC slices from AIR and CIE rats. C) Plot of the relationship between fiber volley amplitude and fEPSP in vHC slices from AIR and CIE rats. D) Representative fEPSPs evoked at 200 µA from AIR and CIE slices. *, p<0.05, ***, p<0.001, AIR =6 slices, 5 rats; CIE n = 8 slices 7 rats).

In the dHC (Fig 2B), there was no main effect of treatment condition on afferent volley amplitude (Figure 4A; F_1, 133_ = 2.112; p = 0.216) but there was a main effect of stimulation intensity on this parameter (F_1, 133_ = 63.662; p <0.001). Although there was no significant interaction between treatment condition and stimulation intensity in the dHC (F_1, 133_ = 1.106; p = 0.44), post hoc analyses revealed a significant increase in afferent volley amplitude in CIE slices at the two highest stimulation intensities, 500 and 700 µA (p = 0.05 at both intensities). Additionally, there was no main effect of treatment condition on fEPSP slope (Figure 4B; F_1, 133_ = 0.862; p = 0.78), but there was a main effect of stimulation intensity on fEPSP slope (F_1, 8_ = 49.184; p<0.001). There was no interaction between treatment condition and stimulation intensity in the dHC (F_1, 8_ = 0.405; p = 0.92) and post hoc analyses confirmed no effect of CIE on fEPSP slope at any of the stimulation intensities tested. When we used a model to conduct an analysis of covariance of all of the dHC data, there was a significant effect of fiber volley amplitude (F_1,166_ = 304.04; p <0.0001), but not CIE, on fEPSP slope (Figure 4C; F_1,166_ = 1.13; p = 0.29), and the interaction between fiber volley amplitude and CIE on fEPSP slope was also significant (F_1,166_=15.35; p <0.0001). This analysis yielded a parameter estimate of -0.567, which indicates that in the dHC of CIE animals, the fiber volley’s positive effect on fEPSP slope is actually decreased, consistent with an overall decrease in excitability.

**Figure 4.**
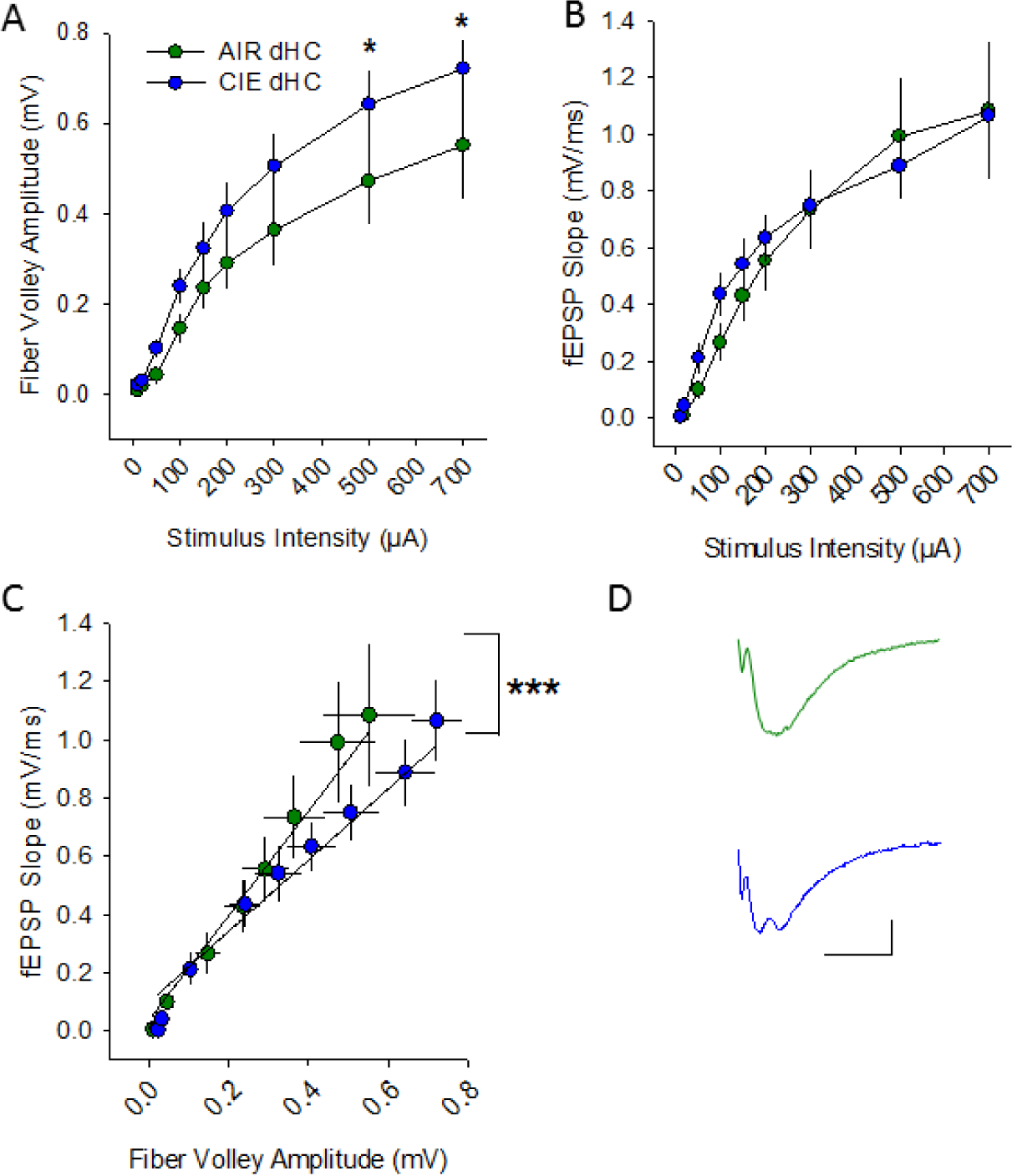
Withdrawal from chronic intermittent ethanol decreases synaptic excitability at dHC Schaffer collateral-CA1 synapses. Plots of stimulation intensity vs fiber volley amplitude (A) and fEPSP slope (B) in dHC slices from AIR and CIE rats. C) Plot of the relationship between fiber volley amplitude and fEPSP in dHC slices from AIR and CIE rats. D) Representative fEPSPs evoked at 200 µA from AIR and CIE slices (*, p<0.05, AIR n = 10 slices, 9 rats; CIE n = 9 slices, 8 rats).

### 3.3. Chronic intermittent ethanol exposure has divergent effects on GluA2 and SK2 protein expression in the ventral hippocampus

In an initial attempt to identify potential mechanisms that may contribute to the marked differences in synaptic adaptation observed in the vHC and dHC following CIE, we used Western blots to analyze protein differences in synaptoneurosome (SN) fractions prepared from the vHC and dHC. These studies focused on the AMPAR GluA2 subunit, whose expression has been linked with CIE-associated increases in synaptic excitability, and the small-conductance Ca^2+^-activated K^+^ channel subunit SK2, a protein that strongly buffers synaptic transmission, particularly in the vHC, and whose expression has been shown to be reduced by CIE (Adelman et al., 2012; Bond et al., 1999; F Woodward Hopf et al., 2010b, 2010a, Mulholland et al., 2011, 2009). We found that, following CIE, there was a significant increase in GluA2 expression in SNs from the vHC, but not dHC, of CIE animals (Figure 5A-D; vHC t(4) = 2.203, p = 0.05, AIR = 1± 0.0298, n = 3; CIE = 1.497 ± 0.224, n = 3; one tailed, unpaired students t-test; dHC t(4) = 1.144, p = 0.11 AIR = 1± 0.088, CIE 1.148± 0.053, n = 3, one tailed unpaired student t-test). Additionally, we found that SK2 expression was decreased in the vHC of CIE animals (Figure 5A&E; t(4) = 3.779, p = 0.01, AIR = 1 ± 0.019, CIE = 0.721± 0.0713, n =3; one tailed, unpaired students t-test), with no changes detected in the dHC (Figure 5B&F; t(4) = 0.994, p = 0.22, AIR = 1± 0.171, CIE = 1.197± 0.993, n = 3; one tailed, unpaired students t-test). Together these findings indicate that following 10 days of CIE and one day of withdrawal, CIE exposure leads to an increase in SN expression of GluA2 and decreases levels of SK2 exclusively in the ventral region of the rat hippocampus.

**Figure 5.**
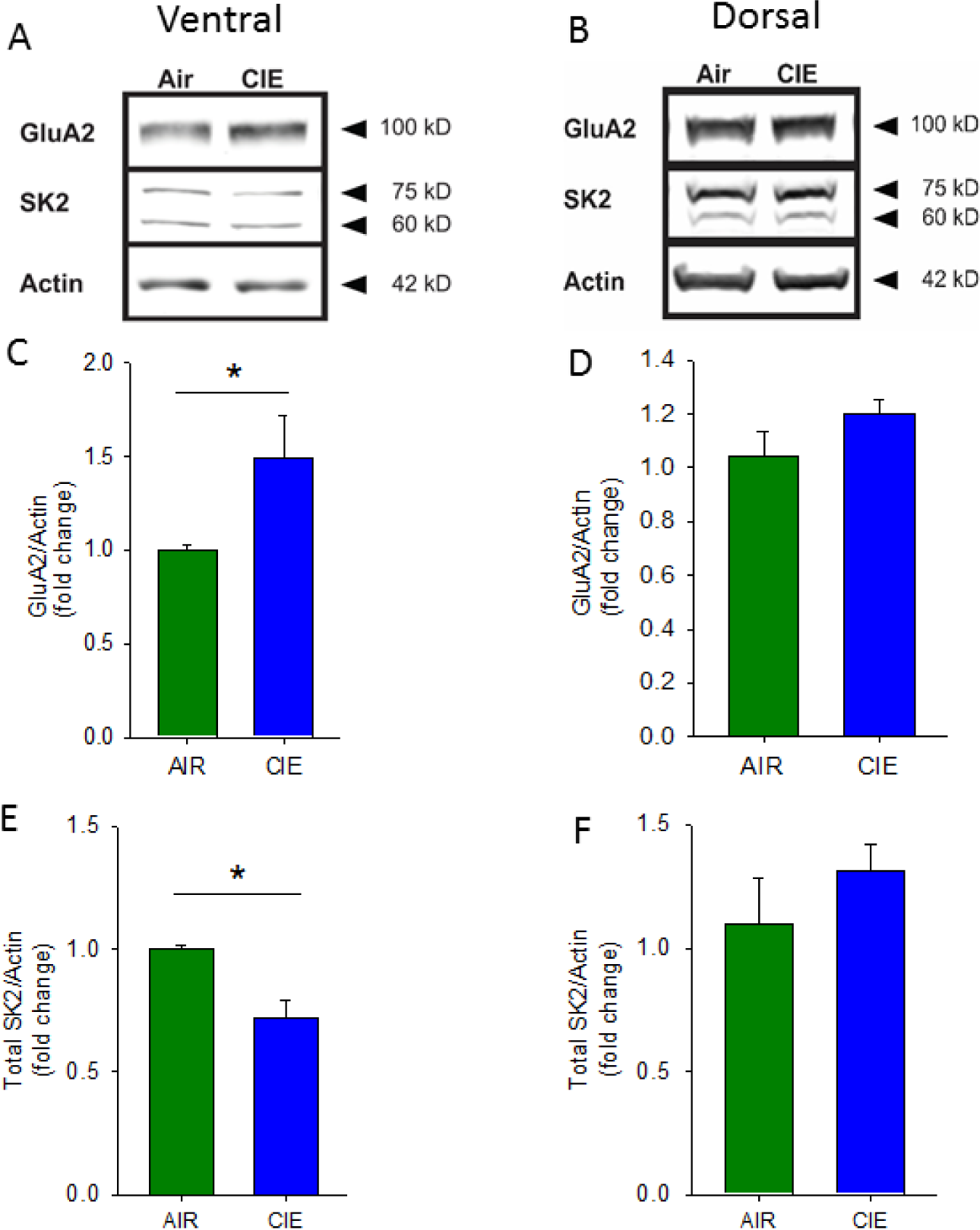
Chronic intermittent ethanol has divergent effects on GluA2 and SK2 protein expression in the ventral and dorsal hippocampus. Representative Western blots of synaptoneurosomes isolated from AIR and CIE vHC (A) and dHC (B) tissue. Group data, normalized to actin, illustrating increased GluA2 protein expression in CIE vHC (C) and not in dHC (D). Group data, normalized to actin, illustrating decreased SK2 protein expression in CIE vHC (E) and not in dHC (F; *, p<0.05, AIR = 4 rats, CIE = 4 rats).

## 4. Discussion

The results of this study demonstrate that withdrawal following CIE produces divergent effects on hippocampal excitability in male Long Evans rats. We first replicated previous work showing that the CIE regimen leads to robust increases in anxiety-like behavior (Läck et al., 2008, 2007, Morales et al., 2018, 2015; Rose et al., 2016). An analysis of the effects of CIE on synaptic excitability within the CA1 region of the vHC and dHC revealed that CIE promoted robust increases in excitability in the ventral subregion but actually led to a significant decrease in overall excitation in the dHC. The CIE-associated increase in synaptic transmission in the vHC was accompanied by a significant increase in SN GluA2 expression and a decrease in SK2 levels. In marked contrast, levels of these synaptic proteins were not affected by CIE in the dHC. Collectively, these studies reveal that CIE-associated increases in synaptic excitability in the hippocampus are exclusively restricted to the ventral domain of this brain region and identify two synaptic proteins that may contribute to the differential effects of CIE on synaptic excitation in the vHC and dHC to CIE.

Withdrawal-induced anxiety symptoms are a hallmark of long term alcohol exposure and play a large role in relapse for individuals with AUD (Driessen, 2001; Schellekens et al., 2015). Not surprisingly, we found that CIE exposure increases anxiety-like behavior on the elevated plus-maze. This finding has been replicated many times in both male and female rodents (Morales et al., 2018, 2015). Moreover, similar CIE effects on anxiety measures have been shown with other behavioral tests. For example, following CIE, C57 mice exhibit an escalation in marble burying, a behavior indicative of an anxiogenic state (Pleil et al., 2015; J. Rose et al., 2016). Using the light/dark box, Lack et al (2007) found that rats spent less time in the light side following CIE exposure, also consistent with an anxiogenic phenotype. Additionally, Morales et al (2018) found that even shorter periods of CIE exposure, as little as 3 days followed by 24 hours of withdrawal, are enough to produce changes in anxiety-like behavior. Additionally, longer periods of CIE exposure (i.e. 2-3 weeks) and withdrawal (i.e. 3 weeks) also results in an increase in anxiety-like behavior on the elevated plus-maze (Valdez et al., 2002). Together, these studies provide strong evidence for the onset and persistence of anxiogensis following CIE exposure.

Several prior studies have employed electrophysiological approaches to characterize the effects of chronic ethanol exposure on measures of hippocampal synaptic activity. For example, Whittington and Little (1990) found that, after 12 weeks of chronic ethanol treatment through voluntary intake and just a few hours of withdrawal (ranging from 3 to 7 hours), hippocampal slices exhibited decreases in the threshold needed to evoke population spikes, indicating an increase in excitability following chronic ethanol. Additionally, intracellular recordings form CA1 pyramidal cells of the hippocampus of rats 3 weeks into withdrawal from a 20 week chronic ethanol liquid diet, showed a significant reduction in inhibitory postsynaptic currents, as well as reduced post-spike afterhyperpolarization when compared with control rats (Durand and Carlen, 1984). Another study, using CIE vapor exposure similar to that employed in this study (2 week of CIE, 24 hr withdrawal), reported increases in synaptic excitability upon increasing stimulation intensities compared with controls (Roberto et al., 2002). This effect was normalized when they examined hippocampal tissue 5 days into withdrawal. Interestingly, using a similar CIE exposure but with no withdrawal (ethanol continuously washed on the slice) or a very brief withdrawal (slices were prepared for recording 2 hours of less after the CIE exposure concluded), the fEPSP responses in the hippocampus were significantly reduced in CIE rats compared to controls (Nelson et al., 1999).

Notably, none of these prior studies differentiated between the ventral and dorsal regions of the hippocampus in their analyses. To that end, we sought to examine the effects of CIE on synaptic excitability within the ventral and dorsal subdivisions of the hippocampus. Analysis of input-ouput curves revealed a significant increase in the amplitude of the presynaptic fiber volley and a marked enhancement of fEPSP slope in recordings from the CA1 region of CIE slices taken from the vHC. A statistical model that was constructed to assess how CIE influenced the interaction between these two synaptic measures revealed a strong overall CIE-dependent increase in synaptic excitability in the vHC. Surprisingly, this same analysis revealed that CIE was associated with a more modest, but significant, decrease in synaptic excitability in the dHC.

These profound subregional differences in synaptic adaptations following CIE are in line with a growing body of evidence showing profound differences in the afferent and efferent circuitry within the vHC and dHC.and the growing appreciation for the differential behavioral roles of these subregions. The vHC playing a critical role in emotional behaviors and the dHC more involved in spatial learning (Michael S Fanselow and Dong, 2010; Henke, 1990; Moser et al., 1995; Moser and Moser, 1998; O’Leary and Cryan, 2014). Lesions of the vHC, but not the dHC, result in decreases in anxiety-like behaviors as well as reduced fear expression (Bannerman et al., 2003, 2004; Kjelstrup et al., 2002). Conversely, lesions of the dHC, but not the vHC, result in impaired learning in the Morris water maze, a well validated test of hippocampus-dependent spatial learning (Moser et al., 1995; O’Leary and Cryan, 2014). Our findings that CIE led to increased anxiety-like behavior and selectively increased synaptic excitability in the vHC are consistent with the critical role of this hippocampal subregion in negative affective behaviors (Moser et al., 1995; Moser and Moser, 1998; O’Leary and Cryan, 2014).

Although a detailed characterization of the mechanisms responsible for the differential responsivity of the vHC and dHC is beyond the scope of this manuscript, our initial biochemical studies identify two synaptic proteins, GluA2 and SK2, whose expression is selectively altered in the vHC, but not dHC, following CIE exposure. The direction of these changes is consistent with an increase in synaptic excitability and interestingly, these proteins have been shown to be altered following CIE in the hippocampus as well as other brain regions (Christian et al., 2012a; Hopf et al., 2010; Mulholland et al., 2011; Durand & Carlen, 1984). One study found that GluA2/3 expression was reduced in the BLA following an identical CIE exposure as this study, indicating that regions that interact with the hippocampus are also exhibiting similar neuroadaptations following this exposure (Christian et al., 2012a). Mulholland and colleagues (2011) found reduced SK2 protein expression in the hippocampus following a modified CIE exposure. Additionally, the calcium-dependent afterhyperpolarization, driven in part by SK channels, was found to be reduced in the hippocampus following three weeks of abstinence from long term ethanol intake (Durand and Carlen, 1984). Both studies did not differentiate between the dHC and vHC but it is possible that the vHC was driving these alterations as the synaptoneurosomal expression of these proteins was not altered by CIE in the dHC. Interestingly, SK channel expression is higher in the vHC than the dHC and these channels buffer glutamatergic synaptic activity to a greater extent in the vHC (Babiec et al., 2017). This differential distribution of SK channels may partly explain the differential effects of CIE on synaptic transmission along the dorsal-ventral axis of the hippocampus

While CIE is a well-validated rodent model of AUD, there are many parallels between the maladaptive behavioral phenotypes promoted by this model and models of stress (Holmes et al., 2012; Morales et al., 2018, 2015). For example, the CIE-induced increases in anxiety-like behavior observed in this study parallel the anxiogenic phenotype that emerges following adolescent social isolation, which has been used to model vulnerability to AUD and anxiety disorders (Almonte et al., 2017; Butler et al., 2016; Rau et al., 2015; Skelly et al., 2015). Notably, both models produce increases in anxiety-like behavior, escalations in voluntary ethanol intake, deficits in fear extinction, and increases in depressive-like behaviors (Holmes et al., 2012; Morales et al., 2018, 2015; Rose et al., 2016; Skelly et al., 2015). Consistent with these behavioral changes, both CIE exposure and adolescent social isolation primarily increase synaptic excitation in the ventral domain of the hippocampus (Almonte et al., 2017). In addition, both CIE and adolescent social isolation increase measures of neuronal excitability in the BLA (Diaz et al., 2011; Läck et al., 2009, 2007; Rau et al., 2015). Notably, we have recently demonstrated that the enhanced intrinsic excitability of BLA pyramidal cells following adolescent social isolation is associated with reduced SK subunit expression (Rau et al., 2015). We further show that administration of a positive SK channel modulator can reduce anxiety-like behavior (Rau et al., 2015). Altogether, these lines of evidence converge upon SK channels as a molecular target that may play an integral role in the neural and behavioral adaptations arising from CIE and adolescent social isolation.

## 5. Conclusions

Together, these findings reveal that CIE-associated increases in hippocampal synaptic excitability are restricted to the ventral domain of this brain region. Given the pivotal role of the vHC in regulating negative affective behaviors, these data may help to explain why increases in anxiety-like behaviors are so prevalent during ethanol withdrawal. The observation that CIE led to a modest decrease in synaptic excitation in the dHC, and that CIE-mediated dysregulation of SN GluA2 and SK2 expression was also restricted to the vHC, add to a growing literature demonstrating that the dHC and vHC are distinct structures that exhibit unique responses to a wide range of environmental manipulations. Given that the intrinsic circuitry of the dHC and vHC is largely preserved but these structures receive largely non-overlapping inputs from different brain areas, it seems likely that unique CIE-associated adaptations upstream of the hippocampus may have contributed to our findings. Future studies, using opto-and chemogenetic circuit mapping techniques, will be needed to identify the afferent inputs to the vHC that promote CIE-associated increases in synaptic excitability in this brain region.

## Acknowledgments

This work was supported by the NIH/NIAAA awards: AA25819 (SEE); AA26117, AA17531, and AA26551 (JLW).

We would also like to thank Kip Zimmerman for his help with the ANCOVA statistical analysis.

